# Medial amygdalar tau is associated with anxiety symptoms in preclinical Alzheimer’s disease

**DOI:** 10.1101/2024.06.03.597160

**Authors:** Joyce S Li, Samantha M Tun, Bronte Ficek-Tani, Wanwan Xu, Selena Wang, Corey L Horien, Takuya Toyonaga, Shreya S Nuli, Caroline J Zeiss, Albert R Powers, Yize Zhao, Elizabeth C Mormino, Carolyn A Fredericks

**Author notes:** Correspondence should be addressed to: Carolyn A. Fredericks, M.D, PO Box 208018, New Haven, CT 06520-8018.

## Abstract

**BACKGROUND:** While the amygdala receives early tau deposition in Alzheimer’s disease (AD) and is involved in social and emotional processing, the relationship between amygdalar tau and early neuropsychiatric symptoms in AD is unknown. We sought to determine whether focal tau binding in the amygdala and abnormal amygdalar connectivity were detectable in a preclinical AD cohort and identify relationships between these and self-reported mood symptoms.

**METHODS:** We examined n=598 individuals (n=347 amyloid-positive (58% female), n=251 amyloid-negative (62% female); subset into tau PET and fMRI cohorts) from the A4 Study. In our tau PET cohort, we used amygdalar segmentations to examine representative nuclei from three functional divisions of the amygdala. We analyzed between-group differences in division-specific tau binding in the amygdala in preclinical AD. We conducted seed-based functional connectivity analyses from each division in the fMRI cohort. Finally, we conducted exploratory post-hoc correlation analyses between neuroimaging biomarkers of interest and anxiety and depression scores.

**RESULTS:** Amyloid-positive individuals demonstrated increased tau binding in medial and lateral amygdala (*F*(4, 442)=14.61, *p*=0.00045; *F*(4, 442)=5.83, *p*=0.024, respectively). Across amygdalar divisions, amyloid-positive individuals had relatively increased regional connectivity from amygdala to other temporal regions, insula, and orbitofrontal cortex. There was an interaction by amyloid group between tau binding in the medial and lateral amygdala and anxiety. Medial amygdala to retrosplenial connectivity negatively correlated with anxiety symptoms (r_s_=-0.103, *p*=0.015).

**CONCLUSIONS:** Our findings suggest that preclinical tau deposition in the amygdala may result in meaningful changes in functional connectivity which may predispose patients to mood symptoms.

## INTRODUCTION

Neuropsychiatric symptoms such as anxiety, irritability, and apathy are both common and debilitating in Alzheimer’s disease (AD). They are the single biggest reason patients must be institutionalized and represent a major source of caregiver distress (1,2). Mood symptoms are early symptoms of AD and can present as early as mid-life. They may, in fact, precede cognitive symptoms (3–5). However, how preclinical Alzheimer’s pathology might give rise to mood symptoms is unclear.

Interestingly, some of the brain regions implicated in emotionality and affect are also among the earliest regions to show AD pathology. For example, the amygdala plays a critical role in emotion modulation and processing (6–9), and pathologic tau deposition occurs in the medial group of the amygdala as early as Braak stage 3 (10), a stage that precedes objective memory impairment for many adults (11). Although early tau deposition in the medial amygdala has been well-established (12,13), research on amygdalar structure and function *in vivo* in AD has been limited. Structural imaging studies have demonstrated low global amygdalar volumes even in the early clinical stages of illness (14,15), and low global amygdalar volume correlates with global illness (16). More recent work has shown that atrophy of the medial and lateral regions of the amygdala predicts conversion from MCI to dementia stage of illness (17). AD patients with neuropsychiatric symptoms including depression, apathy, irritability, and agitation also show differences in functional connectivity of the amygdala when compared with AD patients without affective disturbances (18–21).

Any work that considers the amygdala as a whole is limited by the complex nature of this structure, which is composed of many subnuclei, some of which are heavily impacted by tau pathology (see **Figure 1**, red nuclei), while others are not (**Figure 1**, green nuclei) (12). The most heavily impacted nuclei generally fall within the medial group, including the accessory basal, basomedial, and cortical nuclei and the cortical transition area (10,12,22,23). They are functionally linked, with connectivity to the posterior cingulate cortex, mesial temporal lobe, and ventromedial prefrontal cortex (24). Notably, these structures participate in the default mode network (DMN), which is specifically affected by neurodegeneration in AD and shows decreased connectivity even in very early stage illness (25–29). Conversely, regions spared by AD pathology include the dorsal division nuclei, which participate in an intrinsic connectivity network that is thought to integrate salient socioemotional and homeostatic stimuli, the so-called salience network (SN; see (30) for a concise review), and the lateral division nuclei, which have connectivity predominantly to regions associated with social perception, including fusiform gyrus (31) and superior temporal sulcus (32). The SN is notably spared by AD pathology and shows abnormally increased connectivity in early-stage AD (3,33–35), as does the STS (3,26,36)

**Figure 1.**
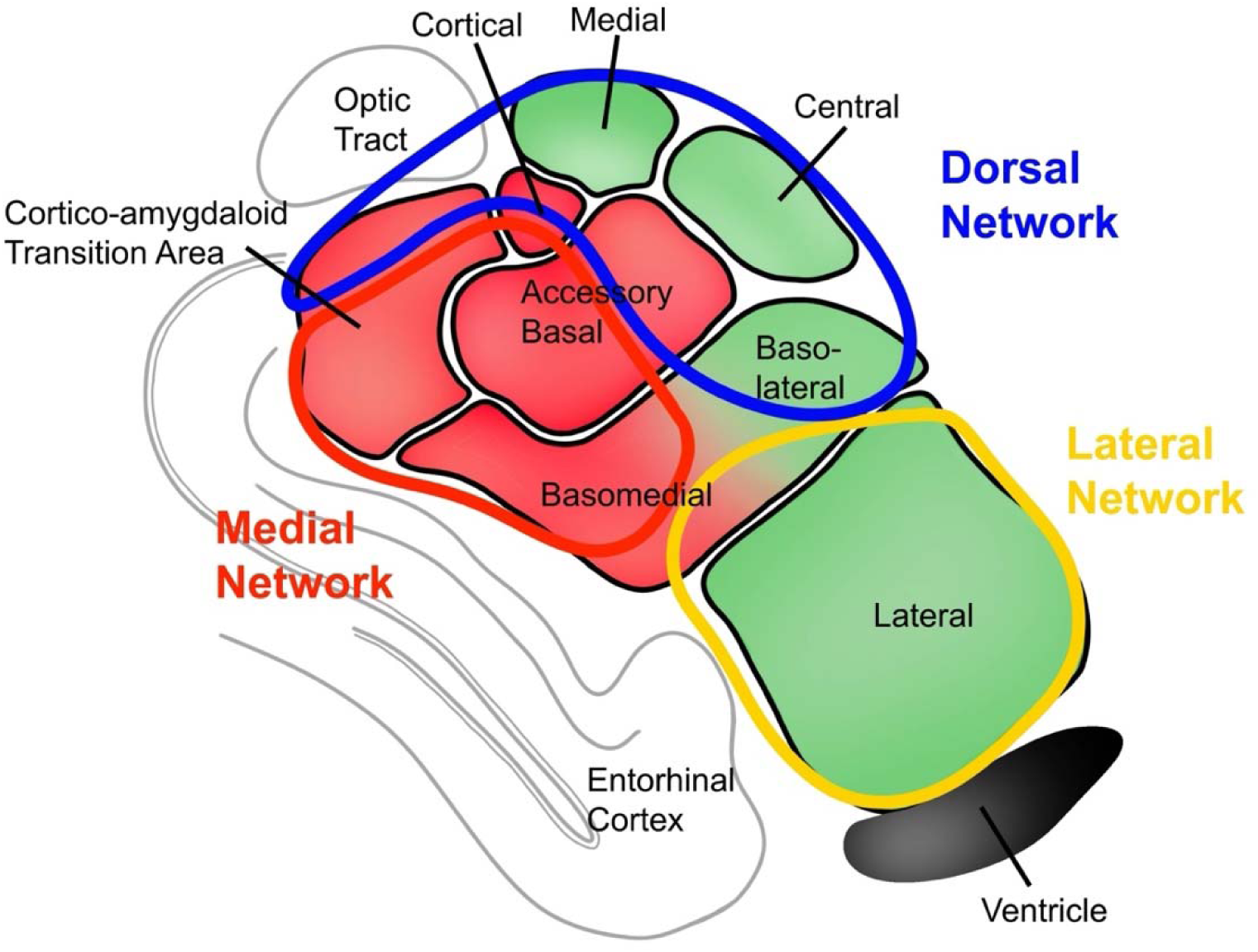
Functional networks overlaid on amygdala tau deposition. Three major functional subdivisions of the amygdala defined by neuroimaging (24) overlaid on a pathologic cross-section through the amygdala showing the major nuclei (based on Saygin et al. 2017 (37)). Red shading indicates nuclei with the heaviest burden of tau pathology (12), while green shading indicates nuclei with relatively less tau burden.

Despite the amygdala’s clear relevance to affective and cognitive circuitry that is markedly impacted in AD, the effects of focal amydalar tau on these networks have not been explored. Preclinical AD populations, in whom early AD pathology (defined by an abnormal burden of amyloid plaques, often accompanied by early-stage neurofibrillary tau tangles) is present in the absence of cognitive symptoms (38), but where early affective symptoms may already be emerging (3–5), offer a particularly meaningful opportunity to assess focal tau signal in the amygdala and related changes in functional circuitry.

Therefore, the purpose of our study was to examine tau PET signal in the amygdala in a large preclinical AD cohort, the Anti-Amyloid Treatment in Asymptomatic AD (A4) Study (39) (n=4486), which includes older individuals with preclinical AD (as defined by positive amyloid PET scan, n=1323), and healthy controls (n=3163). We hypothesized that we could demonstrate focal tau binding in the medial division of the amygdala and that this region would be associated with decreased functional connectivity to DMN regions (specifically posterior cingulate, mesial temporal lobe, and ventromedial prefrontal cortex), while connectivity from other parts of the amygdala might show compensatory increases. On an exploratory basis, we also sought to determine the relationship between tau uptake and abnormal amgydalar connectivity and self-reported mood symptoms.

## METHODS & MATERIALS

### Participants

All analyses used data from the A4 Study (40), a secondary prevention study of amyloid antibody therapy in individuals with preclinical Alzheimer’s disease which has released its baseline (pre-treatment) data to the research community (39). The study recruited thousands of healthy older adults (ages 65-85) who were considered to be at elevated risk for developing Alzheimer’s related memory loss based on age. Participants included in this study (fMRI cohort: n=563; tau PET cohort: n=447 **Table 1**) were cognitively healthy with a Clinical Dementia Rating global score of 0 (41), Mini-Mental State Examination score of 25 or higher (42), and Logical Memory delayed recall score of 6 or higher (43). Of these, n=347 were amyloid-positive (n=251 amyloid-negative), as determined by a standardized uptake value ratio (SUVR) ≥1.15 on an amyloid PET ([^18^F] florbetapir) scan. Institutional review board approval was obtained at each study site and all participants provided written informed consent prior to participation.

**Table 1.** Demographics for A4 Study Data. Bolded text represents the tau PET cohort. Between-group Wilcoxon or chi square tests were performed as appropriate, excluding unknown or not reported values. ^a^ N = 250 for STAI. GDS = Geriatric Depression Scale; STAI = short-form State-Trait Anxiety Inventory.

|  | Amyloid-negative | Amyloid-positive | p-value | **Amyloid-negative** | **Amyloid-positive** | **p-value** |
| --- | --- | --- | --- | --- | --- | --- |
| N | 251 | 312 |  | **100** | **347** |  |
| Age | 70.3 (4.5) | 72.5 (4.9) | <0.001 | **69.8 (4.2)** | **72.4 (4.5)** | **<0.001** |
| Education | 16.5 (2.6) | 16.2 (2.7) | 0.26 | **16.5 (2.8)** | **16.1 (2.8)** | **0.37** |
| Sex (%F) | 62% | 58% | 0.18 | **57%** | **58%** | **0.95** |
| APOE (%E4 carrier) | 19% | 59% | <0.001 | **26%** | **60%** | **<0.001** |
| GDS total | 1.0 (1.4) | 1 (1.3) | 0.32 | **1.1 (1.6)** | **1.0 (1.4)** | **0.65** |
| STAI total | 10.0 (3.3)<sup>a</sup> | 10.2 (3.0) | 0.28 | **10.2 (2.9)** | **10.2 (3.1)** | **0.99** |

### Neuropsychological Measures

All participants underwent a standard neuropsychological battery as well as tests of psychiatric symptomatology, including the Geriatric Depression Scale (GDS) (44) and a six-item short-form of the State-Trait Anxiety Inventory (STAI), focusing on state-anxiety (e.g., “I was worried” on a scale of 1-4) (45). Individuals with significant anxiety or depression were excluded from the study (39). A complete list of tasks is available at ida.loni.usc.edu (“A4 Study Materials”). Participants also underwent functional and structural MRI, amyloid PET ([^18^F]florbetapir), and tau PET (^18^F-AV1451) imaging as detailed below.

### Structural and Functional MRI

Baseline images were downloaded from the A4 LONI Image and Data Archive (v4.1.2021) and are up to date as of October 2023 (39). Participants were scanned on either a 3.0 Tesla Siemens, GE Medical, or Philips Health/Medical Systems scanner. Specific scanning parameters varied by site. T1-weighted structural images were acquired using an MPRAGE sequence (1.05 x 1.05 x 1.2 mm voxel size; matrix size of 240 x 256 x 176 slices; slice thickness=1.2 mm; TR/TI=2300-7636/400-900 ms; TE=2.95-3.11 ms; flip angle=9-11°). Resting-state fMRI sequences were acquired with a 2D gradient-recalled (GR), 2D echo-planar imaging (EPI), or 2D GR EPI sequence (t = 6.5 min; slice thickness=3.3-4mm; TR/TE=3000-3520/30 ms; flip angle=80-90°). Specific acquisition parameters by site can be found at ida.loni.usc.edu (“A4 Study Materials”).

### Structural and functional MR Preprocessing

All structural and functional images were preprocessed using approaches described previously (46,47). Briefly, T1 MPRAGE anatomical scans were skull-stripped through optiBET (48) and registered to an MNI template in BioImage Suite (BIS) (49). In preparation for tau PET analysis, raw T1 MPRAGE anatomical scans also underwent cortical reconstruction and volumetric segmentation of structural MRI images in Freesurfer (www.surfer.nmr.mgh.harvard.edu, version 7.2.0), including skull stripping, subcortical white matter and deep gray matter volumetric segmentation, intensity normalization, surface deformation, and parcellation of gyral and sulcal structures, as a part of its “recon-all” function. Specific details regarding these functions have been described in prior publications (50–56). We then performed high-resolution segmentation of the amygdala and hippocampus using the hippocampal subfields and nuclei of amygdala segmentation package included in Freesurfer 7.2 (37). The Freesurfer amygdala segmentation parcellates the amygdala into the anterior amygdala area, cortico-amygdaloid transition area (CTA), and lateral, basal, paralaminar, accessory basal, medial, central, and cortical nuclei. Of these segmentations, the CTA, central, and lateral nuclei were selected to represent the medial, dorsal, and lateral divisions, respectively.

Functional scans from the A4 dataset were slice-timed and motion-corrected using SPM8. Exclusion criteria for motion were a maximum mean frame-to-frame displacement above 0.3 mm for any scan. Such an approach to handle motion has been shown to limit motion artifact (46,47,57,58). Each participant’s functional scan was registered to their anatomical scan using BIS and smoothed with a 6mm Gaussian kernel in AFNI. Anatomical and functional scans were then resliced and warped in SPM and loaded into CONN toolbox (59) for further preprocessing and seed analysis.

In CONN, anatomical data were segmented into grey matter, white matter, and CSF tissue classes using SPM unified segmentation and normalization algorithm (60,61) with the default IXI-549 tissue probability map template. Denoising of functional data used a standard denoising pipeline in CONN (59), followed by the standard first-level and group-level analyses of seed-based connectivity (see ‘Seed connectivity analyses’ for more). Specific information about the CONN methods for denoising and first- and group-level analyses can be found at https://web.conn-toolbox.org/fmri-methods.

### Amyloid PET

[^18^F]florbetapir PET data was processed by the A4 Study using a previously published pipeline (39). Composite amyloid SUVR was obtained by averaging across six cortical regions from the Automated Anatomical Labeling (AAL) atlas (anterior cingulate, posterior cingulate, lateral parietal, precuneus, lateral temporal, and medial orbital frontal) and normalized to a whole-cerebellum reference region. Amyloid positivity was defined as an SUVR threshold of ≥1.15 or SUVR between 1.10 and 1.15 with a positive visual read, as per A4 Study guidelines (39). The most recent release of amyloid PET SUVR for the A4 dataset resulted in the reclassification of some individuals from positive to negative; we have used the most recent classification here, resulting in an amyloid-negative tau PET cohort of n=100.

### Tau PET

^18^F-AV1451 tau PET scans were downloaded from the A4 LONI Image and Data Archive (v.4.1.2021). Tau PET images were preprocessed and analyzed using PETsurfer (62,63). In brief, five-minute tau PET frames corresponding to 90 to 110 minutes post-injection were motion-corrected and summed. Each participant’s MRI and Freesurfer-generated segmentations were coregistered to the summed PET data. Partial volume correction was applied using the geometric transfer matrix (GTM PVC) method in Freesurfer. Regional SUVR for cortical and amygdalar regions (segmented as described in ‘Structural and functional MR Preprocessing’) in each participant’s T1 MPRAGE coregistered to tau PET were calculated using the whole cerebellum as reference.

### Group differences in tau binding in dorsal and medial amygdala

We used independent t-tests to analyze group differences between whole amygdalar tau binding in amyloid-positive vs amyloid-negative participants. We used ANCOVA to determine whether medial, lateral, and dorsal amygdalar tau SUVR varied by group, using age, intracranial volume (ICV), and entorhinal tau as covariates. We covaried by entorhinal tau because this region has very early tau deposition (Braak stage 1) (10) and is adjacent to parts of the medial and lateral amygdala, where tau signal from the entorhinal cortex could potentially be inaccurately interpreted as regional amygdalar tau due to spatial smoothing. All p-values were corrected for multiple comparisons using Benjamini-Hochberg adjustment (64).

### Seed connectivity analyses

To assess between-group differences in regional amygdalar connectivity, we derived coordinates for the center of mass of the central nucleus (representing the dorsal group), lateral nucleus (representing the lateral group), and CTA (representing the medial group). Due to the concave shape of the CTA, the center of mass was calculated by finding the point within the nucleus with the minimal sum of distances within the area to all the nuclei’s edges. We then derived 2mm radius spherical seeds for the dorsal, lateral, and medial amygdala groups centered at these three coordinates. The averaged fMRI time series was then extracted for each seed ROI and correlated across all voxels across the whole brain. Each subject’s correlation coefficient map was Fisher’s z-to-r transformed, then fit to a second-level regression model for random effects analysis comparing the amyloid-negative and amyloid-positive groups, while controlling for age, sex, handedness, and years of education. All maps were thresholded at a voxelwise threshold of p<0.05 and clusterwise threshold of p<0.05 (FDR corrected). We conducted a conjunction analysis of the three thresholded activation maps from our seed-based analyses by creating an overlay of the three maps using the MarsBaR toolbox in SPM (65). The resulting clusters were then used as clusters of interest in the following correlation analysis.

### Exploratory Correlation Analyses

We conducted exploratory post-hoc analyses to understand the relationship between neuroimaging markers of interest and AD-related neuropsychiatric measures (Geriatric Depression Scale (44) and a short-form of the State-Trait Anxiety Inventory (45)). First, we analyzed associations between tau binding in the three nuclei of interest and mood symptoms using Spearman correlations in each amyloid group. We also assessed if amyloid status was an important interaction term by running a linear regression using the whole sample and including amyloid status as an interaction term in the model. We then analyzed associations between functional connectivity and mood symptoms using partial Spearman correlations to compare mean connectivity values for each cluster (selected from the conjunction analysis of the thresholded activation maps and a cluster of interest from the medial network seed map) and scales of depression and anxiety symptoms, controlling for age, sex, and years of education.

## RESULTS

Participants were well-matched in years of education, sex, and mood symptoms. As anticipated, amyloid-positive individuals were on average older and were more enriched with apolipoprotein E4 (APOE4) genotype (**Table 1**).

### Amyloid-positive individuals showed higher global and medial and lateral amygdalar tau

Consistent with previously published results (66–68), we found significantly greater tau binding in the whole amygdala for amyloid-positive, compared with amyloid-negative, individuals (**Figure 2A**, *t*=8.33 for left amygdala, *p*<0.0001; *t*=9.09 for right amygdala, *p*<0.0001). In the left hemisphere, amyloid-positive individuals had significantly greater binding in medial group (*F*(4, 442)=14.61, *p*=0.00045) and lateral group nuclei (*F*(4, 442)=5.83, *p*=0.024), but no significant difference between cohorts in the dorsal group (*F*(4, 442)=0.17, *p*=0.68), when controlling for age, ICV, and entorhinal tau binding (**Figure 2B**). Similar results were observed in the right hemisphere. Given the importance of tau binding in the entorhinal cortices during the progression of AD, we also conducted ANCOVAs without entorhinal cortex tau as a covariate. In the new models, tau binding in the right central nucleus now significantly differed by amyloid group (*F*(3,443)=5.80, *p=*0.019). Other nuclei were unchanged with regard to significance (**Table S1**).

**Figure 2.**
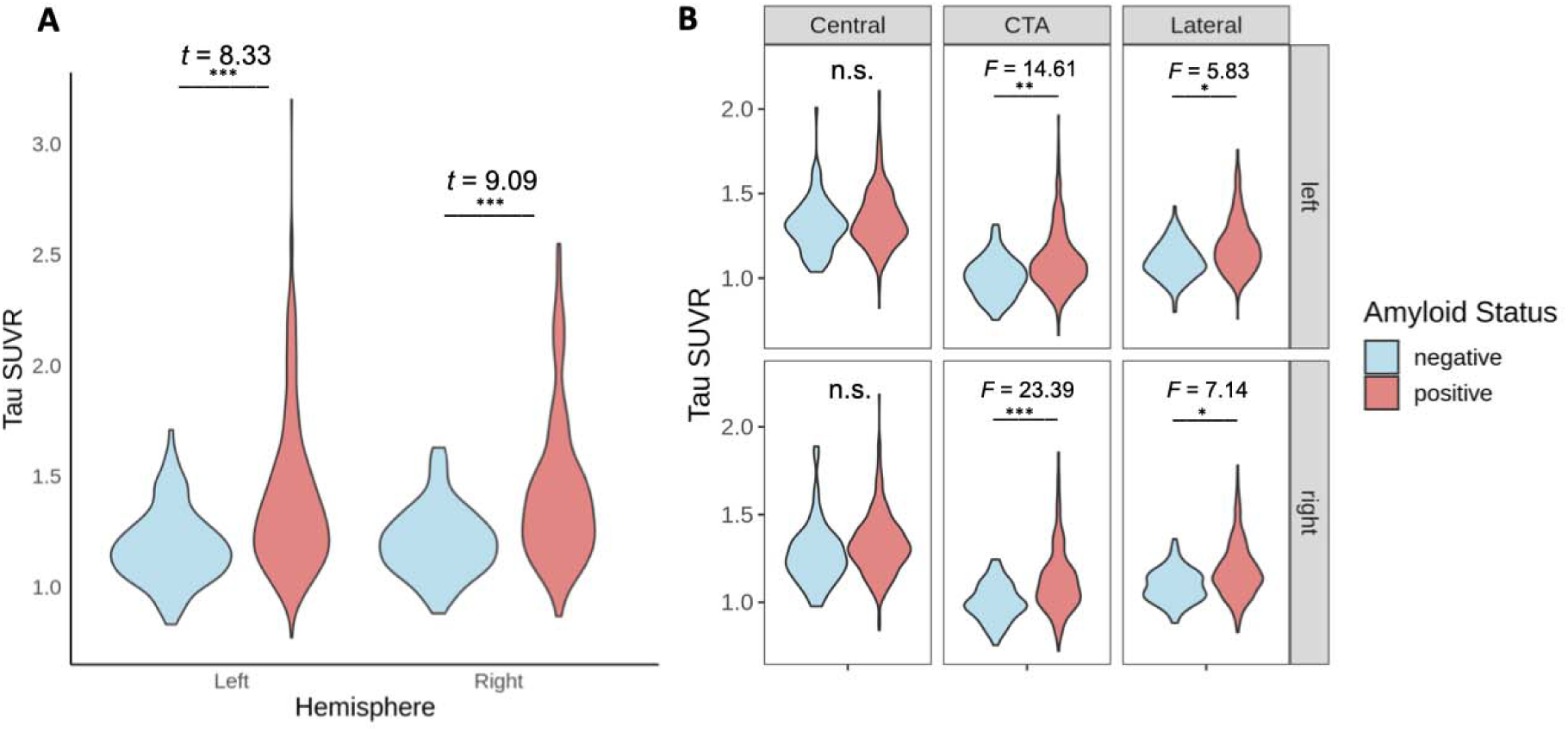
Between-group differences in global and amygdalar region-specific tau binding. (A) Global amygdalar tau binding was greater in the amyloid-positive group. (B) Amygdalar tau uptake was greater in amyloid-positive individuals in the medial (CTA) and lateral (lateral nuclei) amygdala, but not in the dorsal (central nucleus) amygdala. T- or F-statistics reported when appropriate. Results are controlled for age, ICV, and entorhinal tau binding. * p < 0.05, ** p< 0.001, *** p<0.0001. CTA = Cortico-amygdaloid Transition Area; SUVR = Standardized Uptake Value Ratio.

### Amyloid-positive individuals exhibited relatively higher local connectivity to medial and inferotemporal cortex and thalamus, but lower connectivity to insula, accumbens, and retrosplenial cortex

From the dorsal group seed in the central nucleus, amyloid-positive subjects showed elevated connectivity with the bilateral medial temporal lobe, temporal pole and insula relative to amyloid-negative controls (**Figure 3A**). There was decreased connectivity to the basal ganglia, including nucleus accumbens, and left lateral parietal lobe and dorsolateral PFC.

**Figure 3.**
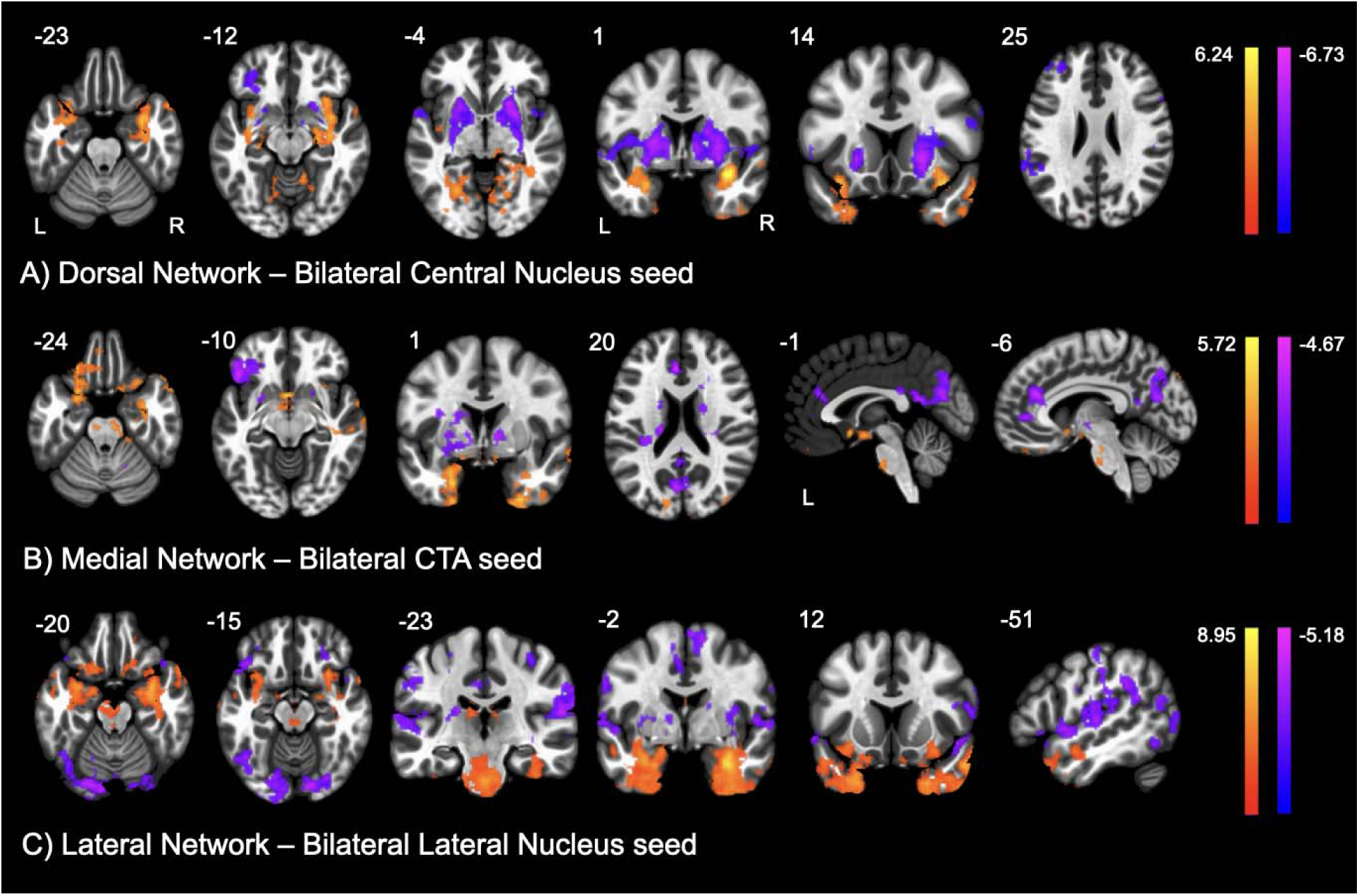
Functional connectivity from amygdalar seeds to the whole brain (amyloid-positive > amyloid-negative control). Across the three seeds, amyloid-positive subjects had greater connectivity from the (A) dorsal network, (B) medial network, and (C) lateral network seeds, to bilateral medial temporal lobes and temporal poles relative to amyloid-negative controls, with variable connectivity to retrosplenial cortex and decreased connectivity to insula, basal ganglia (including accumbens), and superior parietal and dorsolateral prefrontal cortex. Color bars represent T-values. CTA = Cortico-amygdaloid Transition Area.

From the medial group seed in the CTA, amyloid-positive subjects showed higher connectivity to bilateral medial temporal lobe, temporal pole, orbitofrontal cortex, and brainstem, compared to the amyloid-negative group (**Figure 3B**). There was lower connectivity to the retrosplenial cortex (RSC), lateral frontal, and anterior cingulate cortices.

From the lateral group seed in the lateral nucleus, the amyloid-positive group showed increased connectivity to the bilateral medial temporal lobe, temporal pole, orbitofrontal cortex, brainstem, mesial thalamus, and ventral insula (**Figure 3C**). The amyloid-positive group showed decreased connectivity to the superior temporal gyrus, temporoparietal junction, and superior parietal lobe.

Because of its role as a major hub within the DMN (69), we more deeply examined the RSC cluster that demonstrated lower connectivity in amyloid-positive individuals from the medial group seed. In amyloid-negative individuals, average correlated activity from the amygdala to the RSC cluster was highest from the medial group seed and lower from the dorsal and lateral group seeds, in line with our expectation based on the medial group’s connectivity with other DMN structures (24,70). However, in amyloid-positive individuals, activity in the medial seed and RSC was anticorrelated, similar to the other amygdalar seeds, and a significant departure from connectivity in the amyloid-negative state (**Figure 4**).

**Figure 4.**
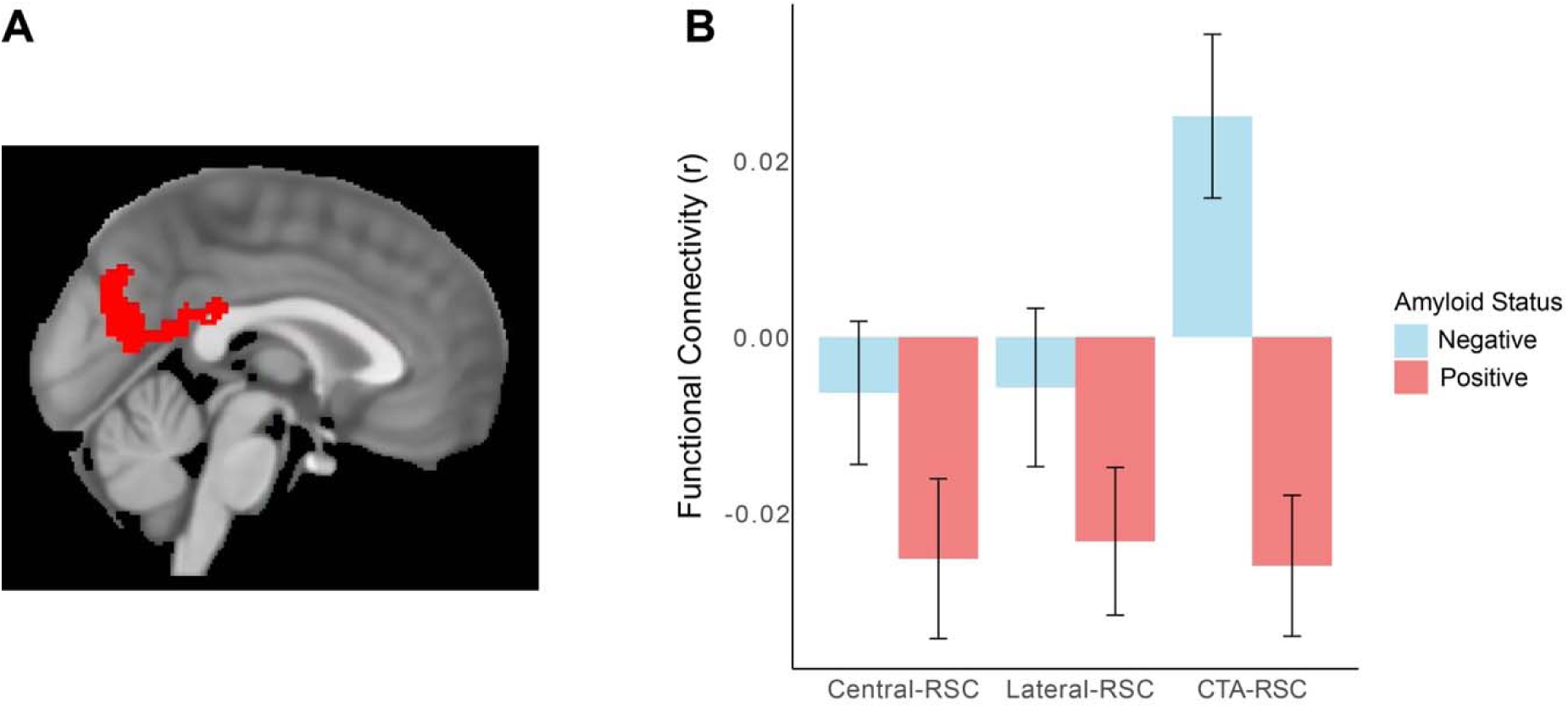
Connectivity to the retrosplenial cortex cluster across the three amygdalar nuclei seeds between amyloid-positive and amyloid-negative groups. (A) Representation of the retrosplenial cluster identified from the CTA seed. (B) Relatively decreased connectivity to the retrosplenial cortex between amyloid groups was only observed from the medial (CTA) seed. Error bars represent standard error of the mean. CTA = Cortico-amygdaloid Transition Area; RSC = Retrosplenial Cortex.

### Tau and connectivity associations with anxiety scores

To assess for a relationship between subclinical mood symptoms and amygdalar tau and connectivity, we conducted exploratory analyses assessing the correlation between (a) focal amygdalar tau signal and (b) amygdalocortical connectivity and neuropsychiatric symptoms (represented by scores on clinical questionnaires for symptoms of depression and anxiety) in preclinical AD. We first examined the relationship between medial-, dorsal-, and lateral-group nuclei tau binding and scores on scales of depression and anxiety (GDS (44) and a short-form state scale of the STAI (45), respectively). There was a significant relationship between lateral division tau uptake and anxiety score in the amyloid-positive group, such that individuals with elevated amyloid had a positive association between tau burden and anxiety score (**Figure 5F**; r_s_=0.12 *p*=0.022). There was a significant interaction by amyloid group in terms of the relationship between depression and tau burden across all three amygdalar nuclei examined, and a significant interaction by amyloid group between anxiety and tau burden in the medial and lateral groups.

**Figure 5.**
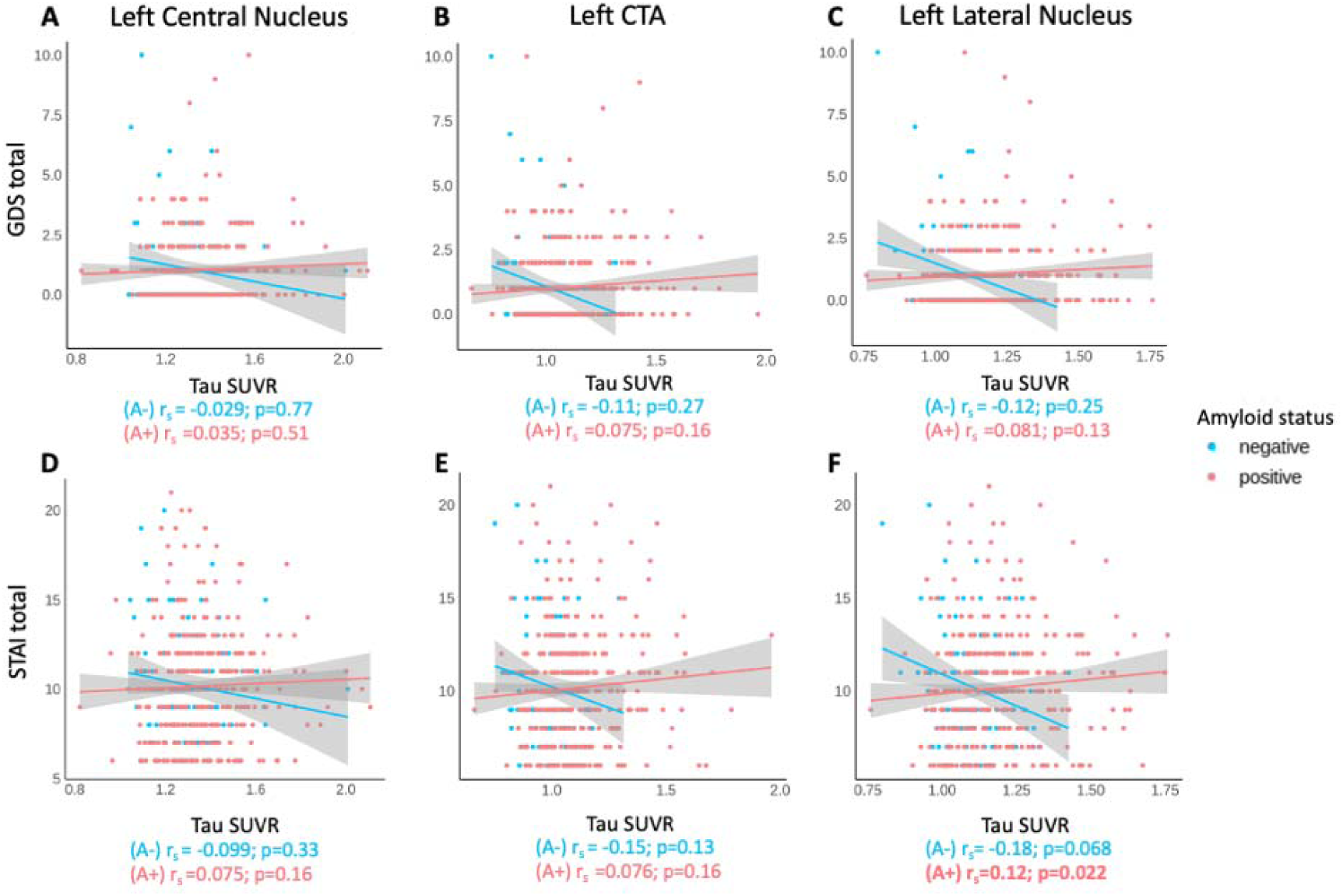
Relationship between medial-, dorsal-, and lateral-group amygdalar tau uptake and depression or anxiety score. Data plotted for amyloid-positive (coral) and amyloid-negative (light blue) individuals. Depression did not significantly associate with (A) dorsal-group (central nucleus), (B) medial-group (CTA), or (C) lateral-group (lateral nucleus) tau binding in the left hemisphere. Anxiety was significantly correlated with (F) left lateral-group tau but was not correlated with (D) dorsal-group or (E) medial-group tau uptake. Shaded area represents 95% confidence. CTA = Cortico-amygdaloid Transition Area; SUVR = Standardized Uptake Value Ratio; GDS = Geriatric Depression Scale score; r_s_ = Spearman Correlation Coefficient; STAI = short-form State-Trait Anxiety Inventory score.

We then assessed for associations between amygdala-to-ROI connectivity and measures of depression and anxiety. Using the intersection (or overlay) of the three seed connectivity maps, we identified three clusters of interest (right hippocampus, left temporal pole, and right insula) which showed consistent differences in connectivity between amyloid-positive and amyloid-negative groups across all three seeds (**Figure 6A**). We also examined the RSC cluster identified only in the medial group (**Figure 4**), given its relevance to DMN function (69). For each of these four clusters and three amygdalar subdivision seeds, we used correlation analyses to assess the relationship between amygdala-to-ROI connectivity and previously described neuropsychiatric tests. Connectivity from the medial group seed to the RSC cluster correlated inversely with anxiety score, such that lower connectivity was associated with higher anxiety (r_s_=-0.103, *p*=0.015, **Figure 6B**).

**Figure 6.**
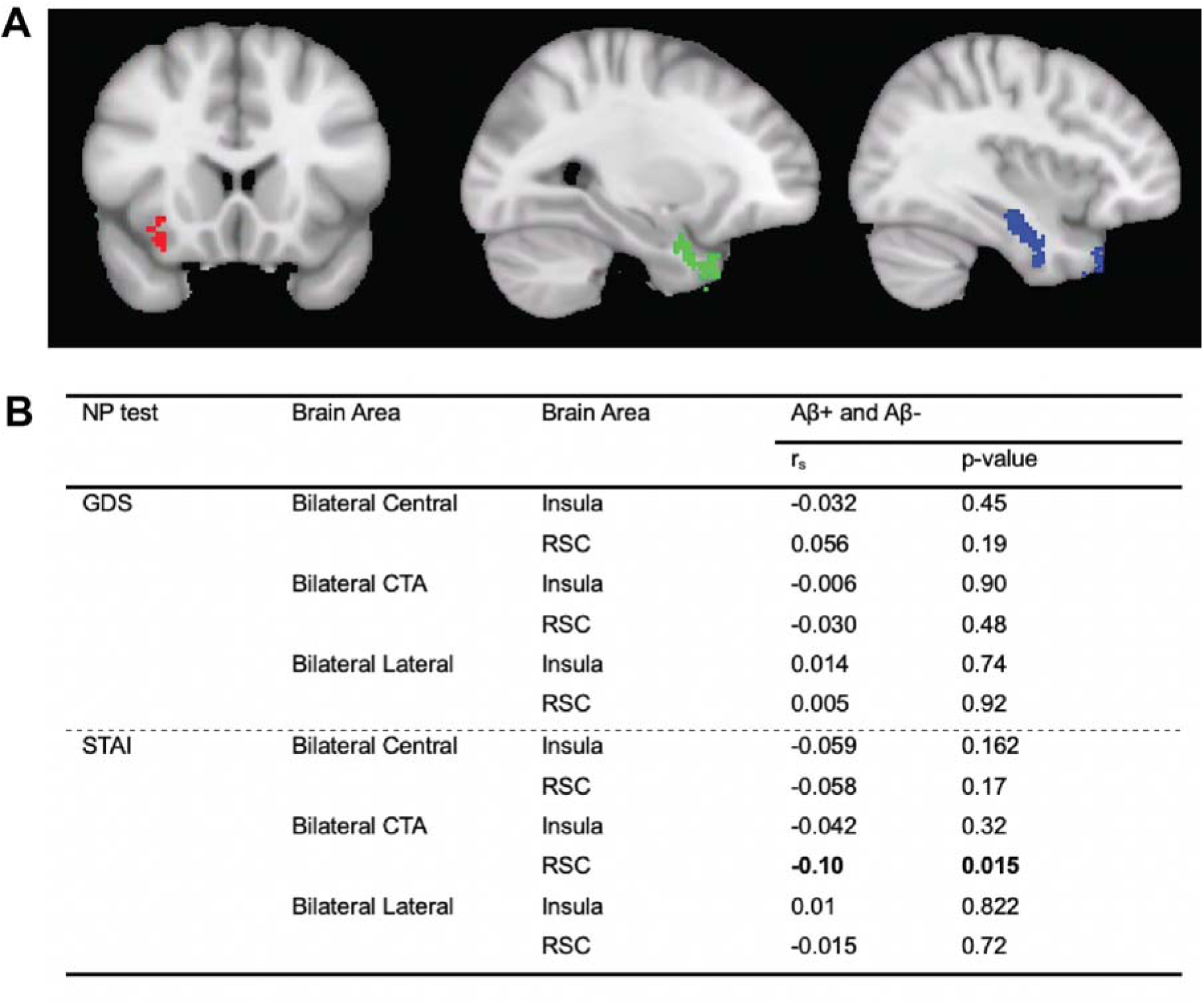
Correlations between clusters of interest and neurocognitive scores. (A) Clusters identified from conjunction analysis of functional connectivity maps from the three amygdalar seeds. Red = right insula cluster; green = left temporal cluster; blue = right temporal cluster. (B) Partial correlation results to neuropsychiatric measures from the dorsal-, medial-, and lateral-groups respectively, to a given cluster. Aβ+ = Amyloid-positive; Aβ-= Amyloid-negative; CTA = Corticoamygdaloid Transition Area; GDS = Geriatric Depression Scale score; NP = Neuropsychiatric Test; RSC = Retrosplenial Cortex; STAI = short-form State-Trait Anxiety Inventory score.

## DISCUSSION

The present study demonstrates that focal increases in tau signal can be detected in pathologically implicated regions in amyloid-positive, cognitively unimpaired individuals, compared to amyloid-negative individuals. Furthermore, we show that these focal increases are associated with changes in functional connectivity and with symptoms of anxiety prior to the emergence of clinically significant cognitive changes. Our results are consistent with post-mortem histological work that shows focal involvement of the amygdala in clinical AD (10,12,13,22,23) and confirm that pathological and functional changes are present in individuals still in the preclinical stages of disease. Our study leverages multimodal techniques and analysis of amygdalar subregions to delineate complex brain-symptom relationships between healthy individuals and those with preclinical AD.

We found significantly greater tau binding in the CTA and lateral nuclei of the amyloid-positive group. This finding generally concurs with previous histology literature which reported greatest tau pathology in the medial regions of the amygdala (10,12,22,23). In addition, our data align with work examining atrophy in the amygdala in MCI converting populations, which found significant atrophy in the right lateral nucleus and CTA, among other medial network nuclei (71). The presence of greater tau binding in the CTA in our preclinical AD cohort agrees with previous literature suggesting tau pathology underlies atrophy in these regions (72). Given the proximity of the ventricles to the central nucleus, our finding of high tau uptake in the central nucleus that did not vary by amyloid status is likely attributable to off-target binding of ^18^F-AV1451 PET ligand to the choroid plexus (73).

Compared with the amyloid-negative cohort, amyloid-positive individuals had elevated connectivity to neighboring limbic regions, including the mesial temporal lobes, temporal poles, insula, and orbitofrontal cortex, a pattern which mirrors increased functional coupling between amygdala and both insula and orbitofrontal cortex in trait anxiety and social anxiety disorder (74–76). Similar findings were seen across all three amygdalar seeds. These findings also align with theories that amyloid positivity promotes local hyperexcitability and increased functional connectivity within the temporal lobe, potentially setting the stage for pathological spread of tau and symptomatic disease (77–80).

All three amygdalar seeds also revealed a pattern in amyloid-positive individuals of relatively depressed connectivity to the basal ganglia, anterior and posterior cingulate, and temporoparietal junction. Interestingly, the medial (CTA nucleus) seed alone revealed a significant difference in functional connectivity between amyloid-negative and amyloid-positive individuals to the RSC, a default mode network hub. While amyloid-negative individuals showed a positive relationship between activity in these two regions, the relationship was inverse in amyloid-positives, suggesting a shift in connectivity between the DMN and the amygdala in preclinical AD.

This shift might relate to early neuropsychiatric symptoms: lower connectivity from medial amygdala to the retrosplenial cluster correlated with higher reported symptoms of anxiety in the A4 cohort. Of all the nodes of the posterior DMN, retrosplenial cortex is perhaps the region with the closest relationship to anxiety: amygdala and RSC play complementary roles in fear extinction (81); the RSC plays a role in fear memory retrieval, and with the amygdala, memory for acute trauma (82,83); and decreased connectivity between the mesial temporal lobe (including the amygdala specifically) and RSC has been implicated in context-dependent fear conditioning in animal studies (84–86). We also observed a positive association between lateral group tau binding and anxiety in the amyloid-positive group, as well as an interaction between amyloid status and mood symptoms in all three amygdalar divisions.

Our study, then, suggests that both the presence of AD pathology in the amygdala, and the shift from a positive to an inverse relationship between medial amygdala and a default mode network hub in amyloid-positive individuals, are linked to higher anxiety scores even in individuals without clinical mood disorder. To our knowledge, our study is the first to demonstrate a significant relationship between mood symptoms and tau in the amygdala in a preclinical AD cohort. This work builds on previous studies that suggest that neuropsychiatric symptoms may begin well before clinically significant cognitive deficits in AD (3–5). Prior studies seeking an association between global amygdalar tau and anxiety or depression did not find one (87,88).

This work has a number of limitations that warrant discussion. The A4 Study had a relatively small sample of amyloid-negative individuals who obtained a tau (^18^F-AV1451) PET scan, limiting our power to examine group differences in tau correlations with mood symptoms. Given off-target binding of the ^18^F-AV1451 ligand to choroid plexus, future work using second-generation tau ligands with different off-target binding profiles will be critical to clarify relationships specifically with the central nucleus, which lies adjacent to the ventricle. In addition, while generally speaking ^18^F-AV1451 binding is specific to tau and not amyloid (89,90), it is possible that the tracer may bind to tau-positive dystrophic neurites within dense-core neuritic plaques; i.e., in this preclinical cohort, some observed signal may be due to amyloid binding and therefore should be interpreted with some caution.

In terms of the relationship between amygdalar tau and mood symptomatology, the A4 Study also excluded individuals with clinically significant anxiety or depression symptoms, potentially eliminating meaningful variation in GDS and short-form STAI scores that would have informed our conclusions. Moreover, this study uses a state-anxiety specific version of the STAI, limiting our ability to generalize our results. Future studies could examine associations in individuals with a broader range of neuropsychiatric scores, including those with clinically significant mood symptoms, who may represent a meaningful and disease-relevant source of variation. Future studies that include more detailed assessment of psychiatric symptomatology (or other metrics assessing emotionality and social function which have been shown to be sensitive to early socioemotional changes in AD (3,91,92)) could better inform our understanding of the earliest affective and behavioral changes associated with AD pathology.

The use of a semi-automated amgydalar segmentation in this study should also be considered. Future studies could more thoroughly test the efficacy of semi-automated vs. manual segmentation efforts, which remains the gold standard. Finally, it will be important to examine longitudinal focal amygdalar tau and functional connectivity over the progression from preclinical to clinical AD to better understand the meaning of these early relationships and how they evolve over the course of illness.

## CONCLUSION

Our results highlight that focal changes in amygdalar tau binding are both present during preclinical AD and associated with anxiety symptoms. Furthermore, amyloid-positivity is associated with a shift in functional connectivity of the amygdala towards higher local temporal connectivity, potentially due to amyloid-mediated hyperconnectivity in temporal circuitry, but lower connectivity to other limbic hubs including insula and accumbens. Notably, inversely correlated activity between the medial amygdala and the retrosplenial cortex, seen only in amyloid-positive individuals, was also associated with symptoms of anxiety. This finding suggests the relationship between medial amygdala and this default mode network hub as a potential focus for future investigations into the emergence of the earliest neuropsychiatric symptoms of AD.

## ACKNOWLEDGMENTS

This work was supported by K23AG059919 (NIH-NIA, CF), T32GM007205 (NIH/NIGMS,CH), an Alzheimer’s Association Clinician Scientist Award (2019-AACSF-644153, CF), the McCance Foundation (CF). CZ is funded through the Yale Alzheimer Disease Research Center (1P30AG066508, PI: Strittmatter, S). AP is supported by a K23 Career Development Award, one R21, and two R01s from the National Institute of Mental Health (K23MH115252-01A1; 5R21MH122940-02; R01MH129721; R01MH131768), by a Career Award for Medical Scientists from the Burroughs-Wellcome Fund, a Carol and Eugene Ludwig Award for Early Career Research, and by the Yale Department of Psychiatry and the Yale School of Medicine.

The A4 Study is a secondary prevention trial in preclinical Alzheimer’s disease, aiming to slow cognitive decline associated with brain amyloid accumulation in clinically normal older individuals. The A4 Study is funded by a public-private-philanthropic partnership, including funding from the National Institutes of Health-National Institute on Aging, Eli Lilly and Company, Alzheimer’s Association, Accelerating Medicines Partnership, GHR Foundation, an anonymous foundation and additional private donors, with in-kind support from Avid and Cogstate. The companion observational Longitudinal Evaluation of Amyloid Risk and Neurodegeneration (LEARN) Study is funded by the Alzheimer’s Association and GHR Foundation. The A4 and LEARN Studies are led by Dr. Reisa Sperling at Brigham and Women’s Hospital, Harvard Medical School and Dr. Paul Aisen at the Alzheimer’s Therapeutic Research Institute (ATRI), University of Southern California. The A4 and LEARN Studies are coordinated by ATRI at the University of Southern California, and the data are made available through the Laboratory for Neuro Imaging at the University of Southern California. The participants screening for the A4 Study provided permission to share their de-identified data in order to advance the quest to find a successful treatment for Alzheimer’s disease. We would like to acknowledge the dedication of all the participants, the site personnel, and all of the partnership team members who continue to make the A4 and LEARN Studies possible. The complete A4 Study Team list is available on: a4study.org/a4-study-team.

