## Supplement Table 1 for "Medial amygdalar tau is associated with anxiety symptoms in preclinical Alzheimer’s disease"

| Hemisphere | Amygdalar nuclei (amygdalar subregion) | *F*-value | *p*-value  (FDR-corrected) |
| --- | --- | --- | --- |
| Left | Central (dorsal) | 2.69 | 0.10 |
|  | **CTA (medial)** | **30.12** | **<0.0001** |
|  | **Lateral (lateral)** | **21.09** | **<0.0001** |
| Right | **Central (dorsal)** | **5.80** | **0.019** |
|  | **CTA (medial)** | **39.86** | **<0.0001** |
|  | **Lateral (lateral)** | **22.22** | **<0.0001** |

**Supplement table 1. ANCOVA analyses examining focal tau uptake between amyloid groups, by amygdalar subregion.**

ANCOVA tests controlled for age and intracranial volume, only. Bolded cells represent amygdalar nuclei that were significant *p* < 0.05. CTA = Cortico-amygdaloid transition area.
